# *Pseudomonas aeruginosa* enhances the efficacy of norfloxacin against *Staphylococcus aureus* biofilms

**DOI:** 10.1101/2020.03.26.010959

**Authors:** Giulia Orazi, Fabrice Jean-Pierre, George A. O’Toole

## Abstract

The thick mucus within the airways of individuals with cystic fibrosis (CF) promotes frequent respiratory infections that are often polymicrobial. *Pseudomonas aeruginosa* and *Staphylococcus aureus* are two of the most prevalent pathogens that cause CF pulmonary infections, and both have been associated with worse lung function. Furthermore, the ability of *P. aeruginosa* and *S. aureus* to form biofilms promotes the establishment of chronic infections that are often difficult to eradicate using antimicrobial agents. In this study, we found that multiple LasR-regulated exoproducts of *P. aeruginosa*, including HQNO, siderophores, phenazines, and rhamnolipids, likely contribute to the ability of *P. aeruginosa* to shift *S. aureus* norfloxacin susceptibility profiles. Here, we observe that exposure to *P. aeruginosa* exoproducts leads to an increase in intracellular norfloxacin accumulation by *S. aureus*. We previously showed that *P. aeruginosa* supernatant dissipates *S. aureus* membrane potential, and furthermore, depletion of the *S. aureus* proton-motive force recapitulates the effect of *P. aeruginosa* supernatant on shifting norfloxacin sensitivity profiles of biofilm-grown *S. aureus*. From these results, we hypothesize that exposure to *P. aeruginosa* exoproducts leads to increased uptake of the drug and/or an impaired ability of *S. aureus* to efflux norfloxacin. Our results illustrate that microbially-derived products can greatly alter the ability of antimicrobial agents to kill bacterial biofilms.

**Importance:** *Pseudomonas aeruginosa* and *Staphylococcus aureus* are frequently co-isolated from multiple infection sites, including the lungs of individuals with cystic fibrosis (CF) and non-healing diabetic foot ulcers. Co-infection with *P. aeruginosa* and *S. aureus* has been shown to produce worse outcomes compared to infection with one organism alone. Furthermore, the ability of these pathogens to form biofilms enables them to cause persistent infection and withstand antimicrobial therapy. In this study, we found that *P. aeruginosa*-secreted products dramatically increase the ability of the antibiotic norfloxacin to kill *S. aureus* biofilms. Understanding how interspecies interactions alter the antibiotic susceptibility of bacterial biofilms may inform treatment decisions and inspire the development of new therapeutic strategies.

## Introduction

Cystic fibrosis (CF) is a genetic, multi-organ disease. Mutations in the cystic fibrosis transmembrane conductance regulator (CFTR) gene result in a defective CFTR channel that causes ionic imbalances leading to thick mucus builds up in the airway, which promotes chronic, polymicrobial infection. Two organisms that are commonly co-isolated from the airways of individuals with CF are *Pseudomonas aeruginosa* and *Staphylococcus aureus* (1). Individually, these pathogens are associated with poor clinical outcomes (2–5); however, it has become clear that co-infection with both *P. aeruginosa* and *S. aureus* can lead to worse prognoses compared to mono-species infections with either of these organisms individually (5–9). In addition, these pathogens frequently co-exist in chronic infections of non-healing wounds, such as diabetic foot ulcers (10, 11).

Both *P. aeruginosa* and *S. aureus* have the ability to form robust biofilms. The biofilm mode of growth enable microbes to survive in the face of extreme environmental challenges, including high doses of antimicrobial agents (12). Moreover, it is becoming increasingly evident that interspecies interactions can have profound consequences on antibiotic susceptibility profiles within polymicrobial biofilm communities (13–19). In particular, multiple studies have uncovered interactions between *P. aeruginosa* and *S. aureus* that influence drug tolerance in diverse ways (13, 17, 18, 20–29).

In this study, we identified an interaction between *P. aeruginosa* and *S. aureus* that potentiates the activity of the nucleic acid synthesis inhibitor norfloxacin against *S. aureus* biofilms. Alone, norfloxacin has a modest impact on *S. aureus* biofilm cell viability; however, we observed a striking 3-4-log increase in the ability of norfloxacin to kill *S. aureus* biofilms in the presence of secreted factors produced by *P. aeruginosa*. Our data are consistent with a model whereby *P. aeruginosa* exoproducts increase intracellular levels of norfloxacin within *S. aureus* cells, leading to increased efficacy of this antibiotic. Thus, our findings could have implications for the treatment of persistent, multi-species infections in the context of chronic pulmonary diseases and non-healing wounds.

## Results

### *P. aeruginosa* supernatant increases *S. aureus* biofilm sensitivity to norfloxacin

Previously, we performed a screen using Biolog Phenotype MicroArray Panels to assess changes in *S. aureus* susceptibility to antibiotics in response to *P. aeruginosa* secreted products (24, 28). This approach allowed us to survey a large diversity of compounds for their antimicrobial activity alone versus in the presence of secreted products from *P. aeruginosa*. In this screen, we found that *P. aeruginosa* cell-free culture supernatant altered the ability of many compounds to kill *S. aureus*, leading to either increased tolerance to some compounds, or alternatively, to increased sensitivity to other compounds. Among the drugs that had increased efficacy against *S. aureus* in the presence of *P. aeruginosa* supernatant were several DNA synthesis inhibitors, including the fluoroquinolone norfloxacin (28). Interestingly, we observed that *P. aeruginosa* exoproducts had the opposite effect on *S. aureus* sensitivity to RNA synthesis inhibitors, whereby *P. aeruginosa* protected *S. aureus* from rifampicin (a rifamycin class antibiotic) and novobiocin (an aminocoumarin class antibiotic) (28).

The Biolog screen allowed us to assess whether combinations of different antimicrobial compounds and secreted products altered the sensitivity of planktonic *S. aureus* cells to these compounds and/or whether these combinations altered the ability of *S. aureus* to form a biofilm. To test whether pre-formed *S. aureus* biofilms exhibited altered sensitivity to norfloxacin in the presence of *P. aeruginosa* exoproducts, we allowed the establishment of *S. aureus* biofilms for 6 h prior to the addition of *P. aeruginosa* supernatant and the antibiotic. Previously, we showed that sessile *S. aureus* cells after 6 h of incubation behave as biofilms under our experimental conditions as judged by their high-level tolerance to multiple antibiotics (24). Additionally, we showed that exposure to *P. aeruginosa* supernatant alone does not alter the cell viability of *S. aureus* biofilms (24, 28).

Here, we found that treatment with *P. aeruginosa* supernatant enhanced norfloxacin’s efficacy against *S. aureus* cells growing as early (6 h) biofilms. Specifically, *P. aeruginosa* supernatant increased the susceptibility of early *S. aureus* biofilms to 12.5 μg/ml of norfloxacin by 3 logs (**Fig. 1A**), and to 25 μg/ml of norfloxacin by 4 logs (**Fig. 1B**). In contrast, *P. aeruginosa* exoproducts did not alter the susceptibility of pre-formed *S. aureus* biofilms to the other DNA synthesis inhibitors tested, including other quinolones (28).

**Figure 1.**
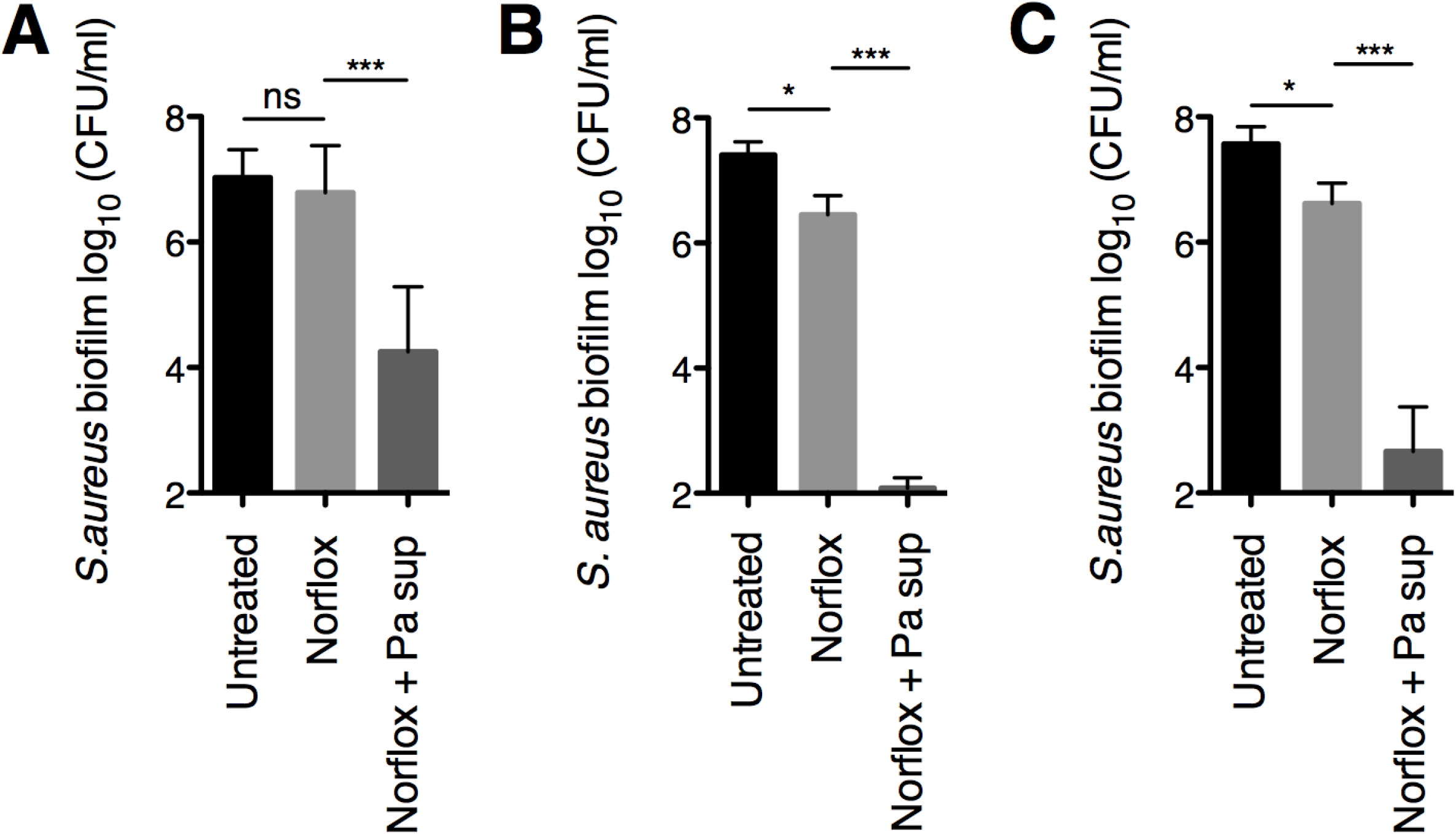
*P. aeruginosa* increases the susceptibility of early and mature *S. aureus* biofilms to norfloxacin. **(A and B)** Biofilm disruption assays on plastic were performed with *S. aureus* Newman, *P. aeruginosa* PA14 supernatant (Pa sup), and norfloxacin (Norflox) at 12.5 μg/ml (**A**) or 25 μg/ml (**B**). Biofilms were grown for 6 hours, and then exposed to the above treatments for 18 hours, and then *S. aureus* biofilm CFU were determined. (**C**) Biofilm disruption assays on plastic were performed with *S. aureus* Newman, *P. aeruginosa* PA14 supernatant (Pa sup), and norfloxacin (Norflox) at 25 μg/ml. Biofilms were grown for 24 hours, exposed to the above treatments for 18 hours, and then *S. aureus* biofilm CFU were determined. Each column displays the average of at least six biological replicates, each with three technical replicates. Error bars indicate SD. ns, not significant; *, p < 0.05, ***, p < 0.001; by ordinary one-way ANOVA and Tukey’s multiple comparison post-test.

Additionally, we tested whether *P. aeruginosa* exoproducts can influence the ability of norfloxacin to kill more mature *S. aureus* biofilms. In this experiment we allowed *S. aureus* biofilms to establish for 24 h prior to treatment with supernatant and the antibiotic. Strikingly, we found that the combination of *P. aeruginosa* supernatant and 25 μg/ml of norfloxacin caused a 4-log reduction in the cell viability of 24-h-grown *S. aureus* biofilms compared to treatment with norfloxacin alone (**Fig. 1C**). Thus, these data indicate that *P. aeruginosa*-produced factors alter the norfloxacin sensitivity profiles of early and more established *S. aureus* biofilms to a similar degree.

### Testing the contribution of *P. aeruginosa* exoproducts to the ability of *P. aeruginosa* to increase *S. aureus* biofilm sensitivity to norfloxacin

Next, we investigated which *P. aeruginosa*-secreted products may be responsible for shifting *S. aureus* biofilm sensitivity to norfloxacin. We first tested the contribution of *P. aeruginosa*-derived factors that were previously found to mediate other *P. aeruginosa-S. aureus* interactions that altered drug sensitivity profiles (13, 24, 25, 28, 30). In particular, we evaluated the contribution of 2-heptyl-4-hydroxyquinoline N-oxide (HQNO) and PQS (*Pseudomonas* quinolone signal), two components of the PQS quorum sensing system biosynthetic pathway, as well as the two siderophores produced by *P. aeruginosa* (pyoverdine and pyochelin), by examining *P. aeruginosa* strains with mutations in these factors (*pqsA*, *pqsH*, *pqsL*, *pvdA*, *pchE*).

We tested the ability of supernatants prepared from the above *P. aeruginosa* strains to alter the susceptibility of *S. aureus* biofilms to norfloxacin at a concentration of 25 μg/ml. We observed that the phenotypes of these individual mutant strains are highly variable (**Fig. 2A**). However, on average we observed that supernatants from strains that are deficient in either the biosynthesis of the entire PQS pathway (Δ*pqsA*), or in the production of HQNO alone (Δ*pqsL*) each had a partial defect in increasing *S. aureus* biofilm sensitivity to norfloxacin relative to wild-type *P. aeruginosa* PA14 supernatant (**Fig. 2A**). Additionally, supernatant from strains lacking either PQS (Δ*pqsH*) or siderophores (Δ*pvdApchE*) had modest defects in shifting drug sensitivity that were not statistically significant. Furthermore, we observed that the supernatant from a *P. aeruginosa* strain lacking the production of HQNO and siderophores (Δ*pqsLpvdApchE*) had a similar phenotype as the Δ*pqsA* and Δ*pqsL* single deletion mutants (**Fig. 2A**).

**Figure 2.**
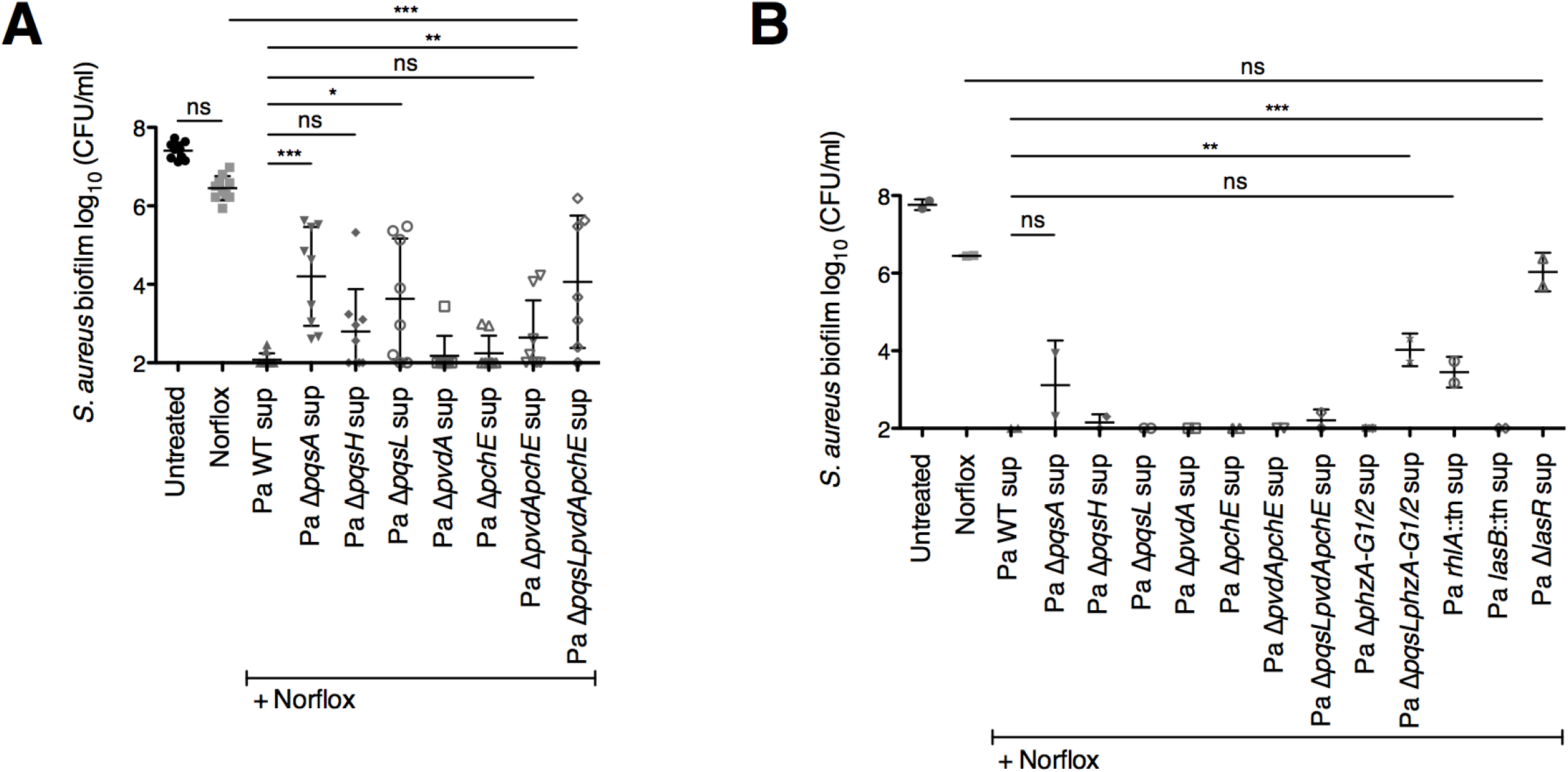
Testing the contribution of *P. aeruginosa*-secreted products to the ability of *P. aeruginosa* PA14 to shift *S. aureus* biofilm sensitivity to norfloxacin. **(A and B)** Biofilm disruption assays on plastic were performed with *S. aureus* Newman, wild-type *P. aeruginosa* PA14 and the specified deletion mutant supernatants (Pa sup), and norfloxacin (Norflox) at 25 μg/ml. Biofilms were grown for 6 hours, exposed to the above treatments for 18 hours, and *S. aureus* biofilm CFU were determined. Each column displays the average of at least seven biological replicates (**A**) or the average of two biological replicates (**B**), each with three technical replicates. Error bars indicate SD. ns, not significant; *, *P* < 0.05, **, p < 0.01, ***, p < 0.001; by ordinary one-way ANOVA and Tukey’s multiple comparison post-test.

Next, we assessed the potential contribution of other *P. aeruginosa*-secreted products to this interspecies interaction. We assayed the following *P. aeruginosa* strains harboring mutations in genes encoding regulatory proteins or genes required for the production of secreted products: the master transcriptional regulator of the Las quorum sensing system (*lasR*), elastase (*lasB*), rhamnolipids (*rhlA*), and phenazines (*phzA-G1/2*) (**Fig. 2B**). Importantly, we previously showed that supernatants from each of the *P. aeruginosa* strains tested here do not alter the viability of *S. aureus* biofilms in the absence of antibiotic (24, 28).

We found that supernatant from a strain that is unable to produce phenazines (Δ*phzA-G1/2*) retained the ability to increase *S. aureus* biofilm sensitivity to norfloxacin (**Fig. 2B**). However, supernatant from a strain that lacks both HQNO and phenazines (Δ*pqsLphzA-G1/2*) had a partial defect in altering *S. aureus* antibiotic susceptibility relative to wild-type *P. aeruginosa* PA14 supernatant. Additionally, supernatant from a strain that is deficient in the production of rhamnolipids (*rhlA*::Tn) led to a modest defect that was not statistically significant. Finally, supernatant from a strain lacking the quorum sensing system transcriptional regulator LasR (Δ*lasR*) completely lacked the ability to enhance the anti-staphylococcal efficacy of norfloxacin (**Fig. 2B**). These results suggest the possibility that HQNO, phenazines, and rhamnolipids – secreted factors that are all under the control of LasR – may play a role in this interspecies interaction.

The combination of *P. aeruginosa* supernatant with norfloxacin at a concentration of 25 μg/ml resulted in *S. aureus* biofilm viability that approaches the limit of detection of our assay, likely leading to the high variability we observed above (**Fig. 2A and B**). Thus, we halved the concentration of norfloxacin used and tested the ability of supernatant from a *P. aeruginosa* strain that cannot produce HQNO and siderophores (Δ*pqsLpvdApchE*) to alter drug sensitivity. At this lower norfloxacin concentration (12.5 μg/ml), we observed that the Δ*pqsLpvdApchE* mutant (denoted Pa ΔΔΔ sup) was completely unable to shift *S. aureus* susceptibility, leading to numbers of viable cells that were not statistically different from treatment with antibiotic alone (**Fig. 3A**). Additionally, we observed much less variability in the response to the combination of norfloxacin and Δ*pqsLpvdApchE* supernatant when using 12.5 μg/ml of drug compared to 25 μg/ml (**Fig. 3A and B**, compare right-most columns). Based on the results from the genetic analyses described above, we hypothesize that multiple LasR-regulated factors (such as HQNO, siderophores, phenazines, and rhamnolipids) may contribute to the ability of *P. aeruginosa* to shift *S. aureus* norfloxacin susceptibility profiles.

**Figure 3.**
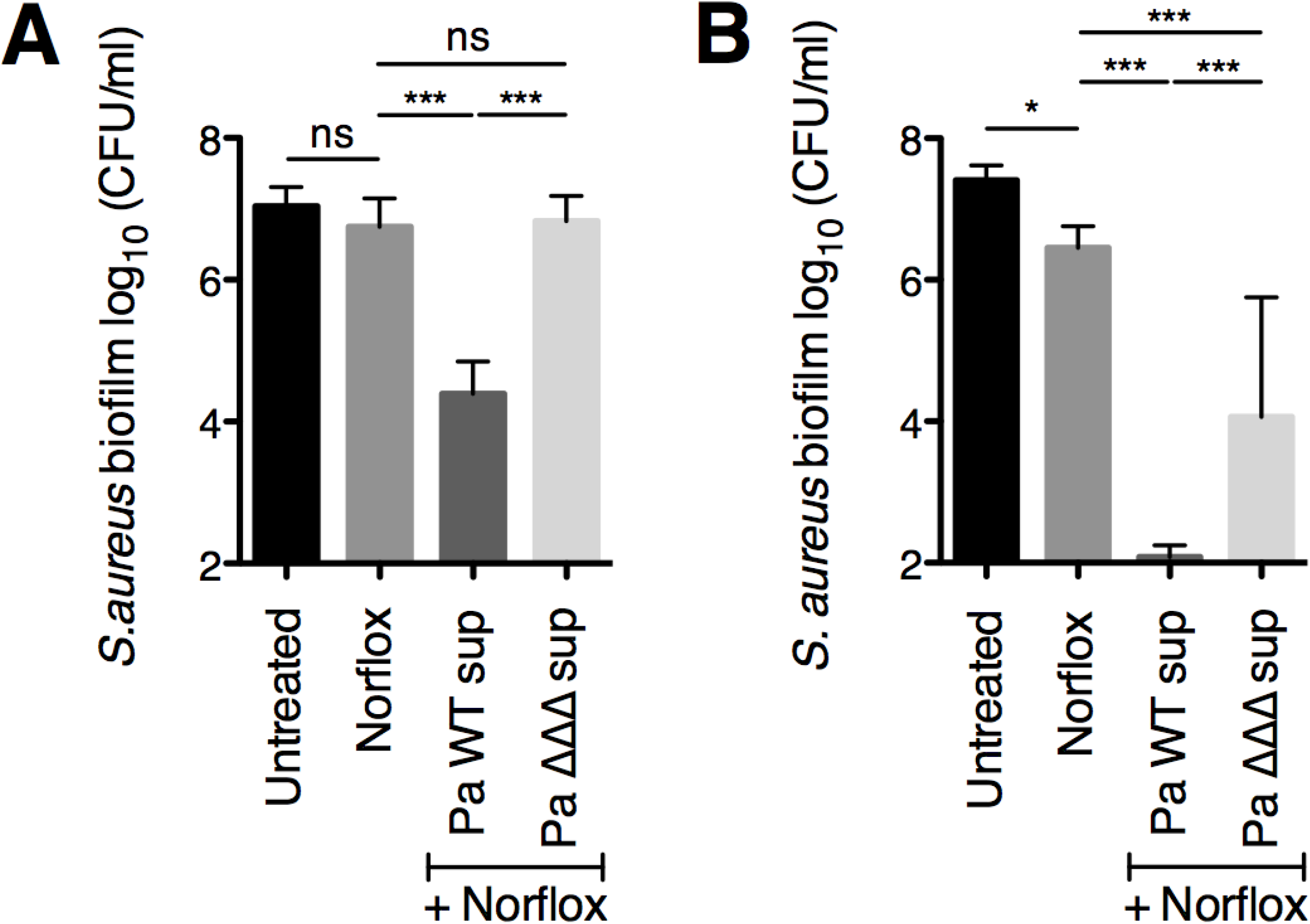
A *P. aeruginosa* strain deficient in the production of HQNO and siderophores is unable to shift *S. aureus* biofilm susceptibility to norfloxacin. **(A and B)** Biofilm disruption assays on plastic were performed with *S. aureus* Newman, supernatants from wild-type *P. aeruginosa* PA14 supernatant (Pa WT sup) and the Δ*pqsLpvdApchE* deletion mutant (Pa ΔΔΔ sup), and norfloxacin (Norflox) at 12.5 μg/ml (**A**) or 25 μg/ml (**B**). Biofilms were grown for 6 hours, exposed to the above treatments for 18 hours, and *S. aureus* biofilm CFU were determined. Each column displays the average of three biological replicates, each with three technical replicates. Error bars indicate SD. ns, not significant; ***, p < 0.001; by ordinary one-way ANOVA and Tukey’s multiple comparison post-test.

### HQNO alone is not sufficient to increase *S. aureus* biofilm sensitivity to norfloxacin

From the results above and our previous studies (24, 28), we hypothesized that the *P. aeruginosa*-produced small molecule HQNO may play a role in increasing the efficacy of norfloxacin against *S. aureus* biofilms. HQNO is a known inhibitor of the *S. aureus* electron transport chain (ETC) (20), and HQNO has been shown to alter *S. aureus* physiology in various ways, including driving *S. aureus* to grow by fermentation (30) and altering membrane fluidity (28). In previous studies performed by our laboratory, we discovered that the addition of pure HQNO alone was sufficient to alter *S. aureus* biofilm susceptibility to the cell wall-targeting antibiotic vancomycin (24), as well as the membrane-active compound chloroxylenol (28). Therefore, we next tested whether the addition of exogenous HQNO alone is sufficient to alter sensitivity to norfloxacin.

The addition of HQNO at a high concentration (100 μg/ml) did not significantly alter the sensitivity of *S. aureus* Newman biofilms to norfloxacin at either concentration tested (12.5 μg/ml or 25 μg/ml norfloxacin) relative to treatment with the antibiotic alone (**Fig. 4A and B**, respectively). This result does not rule out the possibility that HQNO may contribute to the ability of *P. aeruginosa* to alter norfloxacin efficacy; however, these data indicate that HQNO alone is not sufficient for this phenotype.

**Figure 4.**
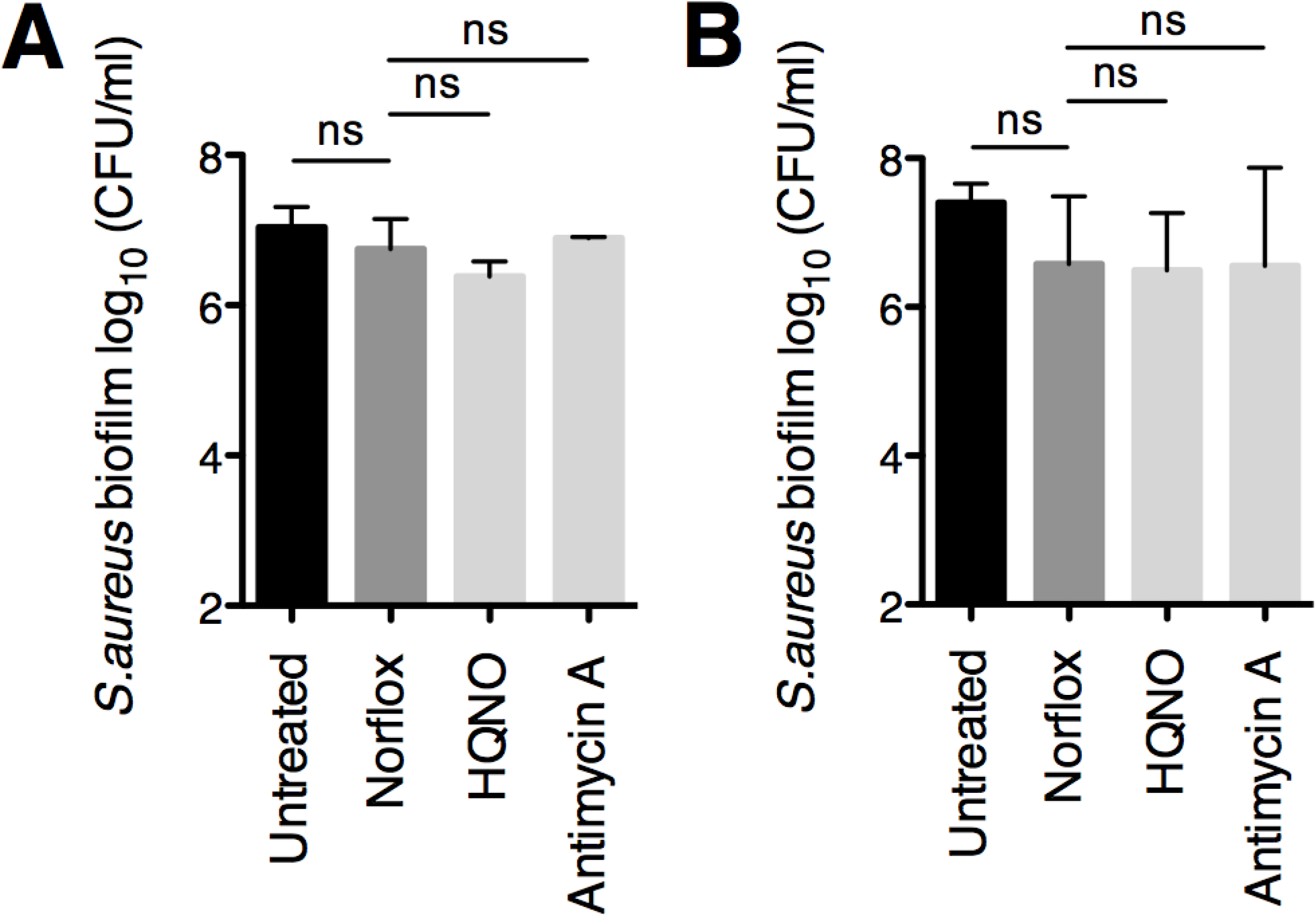
Addition of exogenous HQNO or Antimycin A does not alter *S. aureus* biofilm sensitivity to norfloxacin. **(A and B)** Biofilm disruption assays on plastic were performed with *S. aureus* (Sa) Newman, HQNO at 100 μg/ml, Antimycin A at 100 μg/ml, and norfloxacin (Norflox) at 12.5 μg/ml (**A**) or 25 μg/ml (**B**). Biofilms were grown for 6 hours, exposed to the above treatments for 18 hours, and *S. aureus* biofilm CFU were determined. Each column displays the average of at least two biological replicates, each with three technical replicates. Error bars indicate SD. ns, not significant; by ordinary one-way ANOVA and Tukey’s multiple comparison post-test.

Furthermore, we tested whether treatment with Antimycin A influences *S. aureus* susceptibility to norfloxacin, as this compound is known to inhibit Complex III of the mammalian ETC, but it has been reported to be unable to inhibit the ETC of *S. aureus* (31, 32). Previously, we found that Antimycin A can indeed deplete *S. aureus* membrane potential and that similar to HQNO, Antimycin A increases *S. aureus* biofilm sensitivity to chloroxylenol (28). Here, as in the case of HQNO, we observed that exposure of *S. aureus* biofilms to Antimycin A did not shift susceptibility to norfloxacin at either concentration tested **(Fig. 4A and B**).

### Altering *S. aureus* membrane fluidity influences biofilm sensitivity to norfloxacin

Norfloxacin is believed to enter *S. aureus* cells by diffusion; thus, it is possible that increasing membrane fluidity, which may influence membrane permeability, may facilitate entry of the antibiotic. Our findings from a previous study suggest that HQNO has a fluidizing effect on the *S. aureus* cell membrane (28). Here, we tested whether altering membrane fluidity can impact *S. aureus* sensitivity to norfloxacin.

To test this hypothesis, we first treated *S. aureus* Newman biofilms with the known fluidizing agent benzyl alcohol (33–35) in combination with norfloxacin. We observed an approximate 1-log reduction in *S. aureus* Newman biofilm cell viability upon exposure to the combination of benzyl alcohol and norfloxacin relative to treatment with norfloxacin alone (**Fig. 5A**). In addition, we tested whether the opposite change – decreased fluidity – impacts *S. aureus* susceptibility to this antibiotic. Upon addition of DMSO, a compound that has been described to increase the rigidity of cell membranes (36, 37), we did not observe any changes in antibiotic sensitivity (**Fig. 5B**).

**Figure 5.**
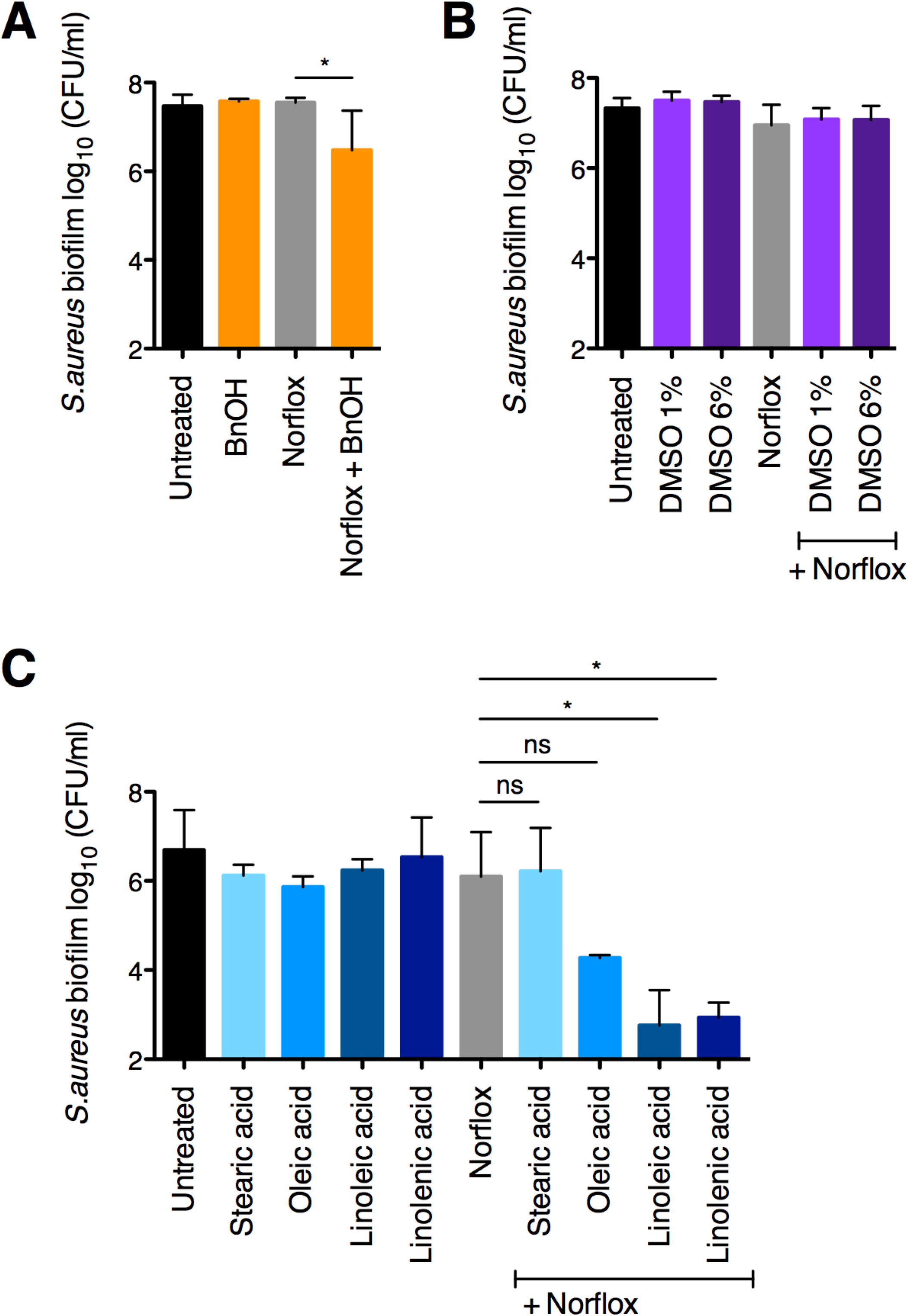
Testing the effect of changing membrane fluidity on the efficacy of norfloxacin against *S. aureus* biofilms. (**A and B**) Biofilm disruption assays on plastic were performed with *S. aureus* Newman, norfloxacin (Norflox) at 25 μg/ml, benzyl alcohol (BnOH) at 50 mM (**A**), and dimethyl sulfoxide (DMSO) at 1% and 6% (**B**). Biofilms were grown for 6 hours, exposed to the above treatments for 18 hours, and *S. aureus* biofilm CFU were determined. Each column displays the average from at least three biological replicates, each with three technical replicates. (C) Biofilm disruption assays on plastic were performed with *S. aureus* Newman, norfloxacin (Norflox) at 12.5 μg/ml, and the specified fatty acids at 100 μg/ml. Biofilms were grown for 6 hours, exposed to the above treatments for 18 hours, and *S. aureus* biofilm CFU were determined. Each column displays the average from two biological replicates, each with three technical replicates. Error bars indicate SD. ns, not significant; *, *P* < 0.05; by ordinary one-way ANOVA and Tukey’s multiple comparison post-test.

*S. aureus* is known to incorporate unsaturated fatty acids into the cell membrane (38, 39), which has been shown to increase membrane fluidity (39, 40). Here, we observe that the addition of unsaturated fatty acids increased *S. aureus* biofilm sensitivity to norfloxacin by 2-3 logs relative to treatment with the drug alone (**Fig. 5C**). Therefore, these data suggest that modulating *S. aureus* membrane fluidity has the potential to influence the efficacy of norfloxacin.

It is possible that by increasing membrane fluidity, the above treatments also change the permeability of the membrane, which may allow for increased entry of norfloxacin into the cell and explain the enhancement of antibiotic-mediated killing we observed. The findings that HQNO enhances membrane fluidity (28), but does not increase sensitivity to norfloxacin on its own **(Fig. 4**) indicate that the HQNO-mediated change in membrane fluidity we observed is not sufficient to explain our findings in this study.

### Exposure to *P. aeruginosa* exoproducts increases the intracellular accumulation of norfloxacin by *S. aureus*

As described above, a potential mechanism by which *P. aeruginosa* potentiates the anti-staphylococcal efficacy of norfloxacin is by causing an increase in *S. aureus* intracellular levels of this antibiotic. Intracellular drug accumulation can be influenced by changes in uptake as well as efflux. In *S. aureus*, norfloxacin uptake is thought to occur via simple diffusion (41–43). In contrast, norfloxacin efflux is mediated by multidrug efflux pumps that are powered by the proton-motive force (PMF) (44). Thus, norfloxacin efflux can be inhibited via the addition of the proton ionophore carbonyl cyanide 3-chlorophenylhydrazone (CCCP) (45).

Here, we tested whether *P. aeruginosa*-secreted factors alter the intracellular accumulation of norfloxacin by *S. aureus*. In this experiment, we assessed whether norfloxacin internalization by *S. aureus* would be altered in the presence of *P. aeruginosa* supernatant by taking advantage of norfloxacin’s intrinsic fluorescence. As a control, we showed that exposure of the cells to CCCP at a concentration of 5 μM led to a significant increase in *S. aureus* intracellular levels of norfloxacin relative to exposure to drug alone (**Fig. 6A**).

**Figure 6.**
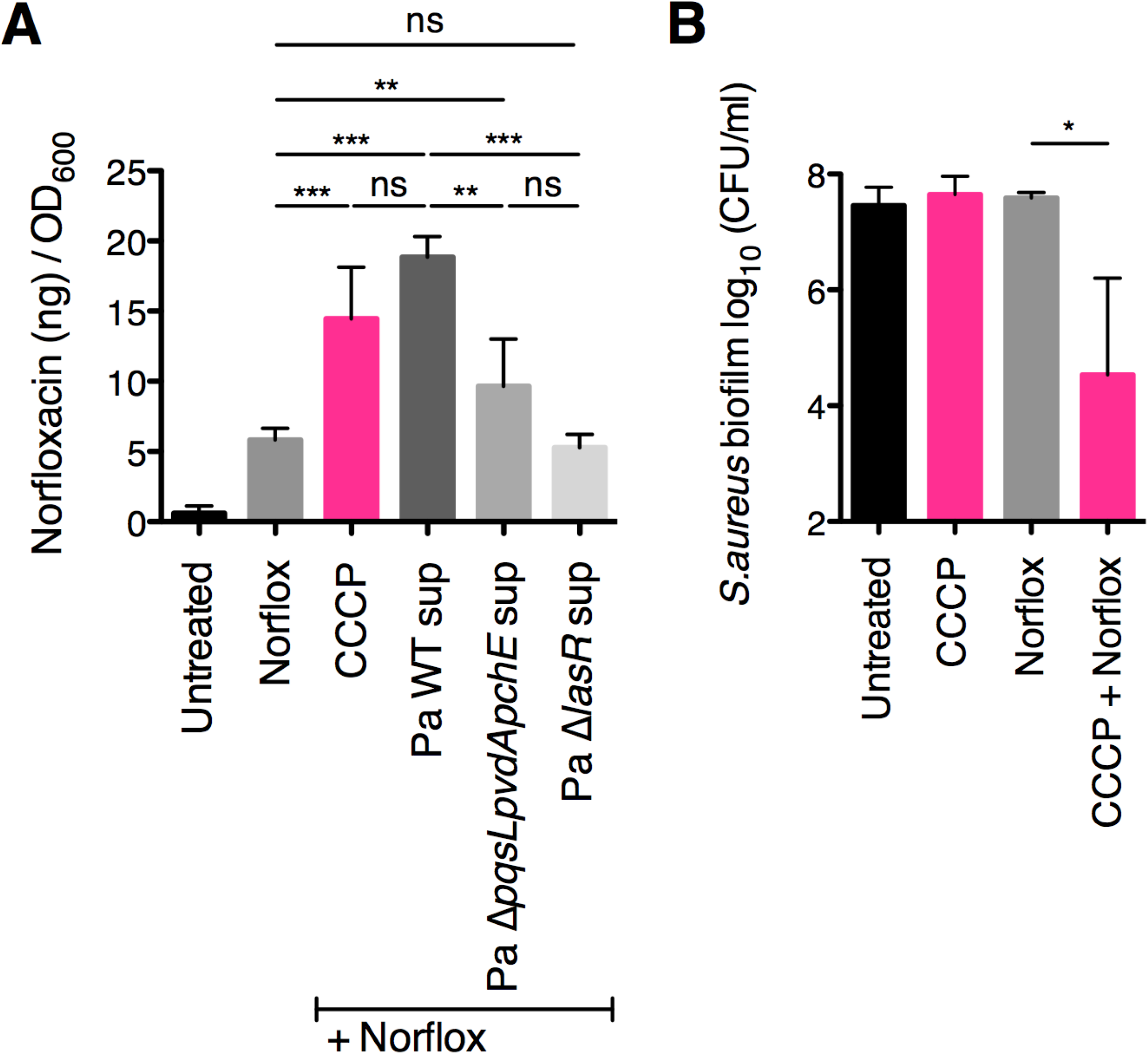
Exposure to *P. aeruginosa* exoproducts increases *S. aureus* intracellular norfloxacin levels. (**A**) Antibiotic internalization assays were performed with *S. aureus* Newman, wild-type *P. aeruginosa* PA14 and the specified deletion mutant supernatants (Pa sup), carbonyl cyanide 3-chlorophenylhydrazone (CCCP) at 5 μM, and norfloxacin (Norflox) at 10 μg/ml. Intracellular norfloxacin levels were quantified by measuring the intrinsic fluorescence at 440 nm following excitation at 281 nm, and a standard curve was used to relate fluorescence values to the amount of intracellular norfloxacin (μg) in each sample. Values are reported as the amount of intracellular norfloxacin (μg) / OD_600_. Each column displays the average of three biological replicates, each with three technical replicates. Error bars indicate SD. ns, not significant; **, p < 0.01, ***, p < 0.001; by ordinary one-way ANOVA and Tukey’s multiple comparison post-test. (**B**) Biofilm disruption assays on plastic were performed with *S. aureus* Newman, CCCP at 5 μM and norfloxacin (Norflox) at 25 μg/ml. Biofilms were grown for 6 hours, exposed to the above treatments for 18 hours, and *S. aureus* biofilm CFU were determined. Each column displays the average of three biological replicates, each with three technical replicates. Error bars indicate SD. ns, not significant; *, *P* < 0.05; by ordinary one-way ANOVA and Tukey’s multiple comparison post-test.

Interestingly, we found that exposure to wild-type *P. aeruginosa* supernatant (Pa WT sup) led to *S. aureus* intracellular levels of norfloxacin that were significantly higher than exposure to norfloxacin alone, and were not significantly different from treatment with CCCP (**Fig. 6A**). Additionally, we observed that exposure to supernatant from the *P. aeruginosa* Δ*pqsLpvdApchE* deletion mutant resulted in significantly lower *S. aureus* intracellular norfloxacin levels relative to treatment with wild-type *P. aeruginosa* supernatant (**Fig. 6A**). Furthermore, treatment with supernatant from the *P. aeruginosa* Δ*lasR* deletion mutant entirely abrogated the ability of *P. aeruginosa* supernatant to increase *S. aureus* norfloxacin accumulation, leading to intracellular drug levels that were not significantly different from exposure to the drug alone (**Fig. 6A**). Together, these data suggest that treatment with *P. aeruginosa* exoproducts increases *S. aureus* intracellular accumulation of norfloxacin, which may occur via modulation of influx and/or efflux of this antibiotic.

Furthermore, the addition of CCCP at 5 μM significantly enhanced the anti-staphylococcal efficacy of norfloxacin at 25 μg/ml under our study conditions (**Fig. 6B**). Together, these data suggest that a decrease in PMF can potentiate the activity of norfloxacin against *S. aureus* biofilms perhaps via allowing increased intracellular accumulation of this antibiotic.

### *P. aeruginosa* supernatant does not impact the expression of selected *S. aureus* antibiotic transporters

Previously, our laboratory performed an RNA-seq experiment to determine whether *S. aureus* gene expression changes during co-culture with *P. aeruginosa* (30). We observed that multiple *S. aureus* drug transporters are differentially regulated in the presence of *P. aeruginosa* (these data are summarized in **Table 1**). However, the following caveats to this experiment should be considered: *S. aureus* was co-cultured with *P. aeruginosa* instead of being exposed to *P. aeruginosa* cell-free culture supernatants, and the RNA-seq experiment was performed on a single biological replicate.

**Table 1.** Testing strains with transposon insertions in genes coding for *S. aureus* antibiotic transporters that were differentially regulated upon co-culture with *P.* aeruginosa cells^a^.

| Gene ID<sup>b</sup> | Tn ID<sup>c</sup> | Gene name | Gene Annotation | Fold change<sup>d</sup> |
| --- | --- | --- | --- | --- |
| 02418 | NE12 | *lmrS* | drug resistance transporter,<br>EmrB/QacA subfamily | - 6.7 |
| 01876 | NE17 |  | putative drug transporter | - 3.7 |
| 02700 | NE179 | *mdeA* * | multidrug resistance protein | - 14.9 |
| 02003 | NE370 |  | toxin exporting ABC transporter,<br>permease-ATP-binding protein,<br>putative | - 11.4 |
| 00315 | NE405 | *mepA* * | MATE efflux family protein | - 7.8 |
| 02420 | NE531 | *sdrM* * | putative drug transporter | - 2.9 |
| 00078 | NE749 | *sbnD* | putative drug transporter | + 1.3 |
| 02818 | NE773 |  | drug transporter | - 6.0 |
| 02629 | NE781 |  | multidrug resistance protein B, drug<br>resistance transporter | - 5.1 |
| 00246 | NE901 |  | putative drug transporter | - 1.8 |
| 00703 | NE1034 | *norA* * | multidrug resistance protein | - 1.6 |
| 02531 | NE1041 |  | putative drug transporter | - 9.8 |
| 01448 | NE1184 | *norB* * | putative drug transporter | + 11.7 |
| 02740 | NE1400 |  | putative drug transporter | - 4.1 |
| 00058 | NE1804 | *norC* * | putative drug transporter | - 2.0 |
| 02419 | NE1616 | *sepA* | hypothetical protein | - 11.8 |

<sup>a</sup> Reported are data from an RNA-seq experiment from a previous study from our laboratory (30).
Briefly, *S. aureus* 8325-4 was co-cultured with *P. aeruginosa* PA14 cells or grown in mono-
culture in MEM + L-Gln on CF-derived epithelial cells for 6 h prior to RNA isolation and
sequencing. The fold change shown indicates the expression in co-culture versus the mono-
culture of *S. aureus*, with negative numbers indicating lower expression in co-culture. Note:
Only a single biological replicate was performed for this experiment.
<sup>b</sup> SAOUHSC gene number.
<sup>c</sup> Transposon identification number.
<sup>d</sup> co-culture/mono-culture from a single replicate RNA-seq experiment.
\*genes coding for antibiotic transporters that are reported to efflux norfloxacin.

Norfloxacin is a substrate of several multidrug efflux pumps belonging to the major facilitator superfamily (MFS) (NorA, NorB, NorC, MdeA, and SdrM), as well as the multidrug and toxic compound extrusion (MATE) family member MepA (44, 46).

Among these genes, only *norB* expression increased during co-culture (11.7-fold increase in co-culture, **Table 1**). In contrast, all of the other known norfloxacin transporters were downregulated in the presence of *P. aeruginosa* cells. The expression of *norA*, *norC*, and *sdrM* was minimally altered during co-culture (1.6-fold decrease, 2-fold decrease, and 2.9-fold decrease, respectively), whereas *mdeA* and *mepA* were strongly downregulated in the presence of *P. aeruginosa* cells (14.9-fold decrease and 7.8-fold decrease, respectively, **Table 1**).

Moreover, the expression of two genes encoding putative drug transporters also decreased in the presence of *P. aeruginosa* cells: SAOUHSC_02003 decreased 11.4-fold in co-culture; SAOUHSC_02531 decreased 9.8-fold in co-culture (**Table 1**). Similar changes in the expression of these two genes were observed in other studies that examined the transcriptional response of *S. aureus* to either co-culture with *P. aeruginosa* (47), or exposure to anoxic conditions (48).

From the results above, we hypothesized that *P. aeruginosa* exoproducts decrease the expression of multiple *S. aureus* drug efflux pumps. To test this hypothesis, we performed qRT-PCR to evaluate the effect of *P. aeruginosa* supernatant on the expression of *S. aureus* drug transporters. We selected the following subset of genes: *mdeA*, *mepA*, *sdrM, norA*, based on the RNA-seq results above and their known role in norfloxacin efflux.

Exposure to *P. aeruginosa* supernatants (prepared either from the wild-type or the Δ*lasR* mutant) did not lead to significant changes in expression of the four drug transporters above relative to exposure to medium alone (**Fig. 7**). These results suggest that *P. aeruginosa* exoproducts have minimal impact on the expression of at least a subset of *S. aureus* antibiotic transporters under our study conditions. Therefore, our data do not support a model whereby downregulation of *S. aureus* efflux pumps contributes to the *P. aeruginosa*-mediated increase in *S. aureus* intracellular norfloxacin levels and reduced biofilm cell viability we observe.

**Figure 7.**
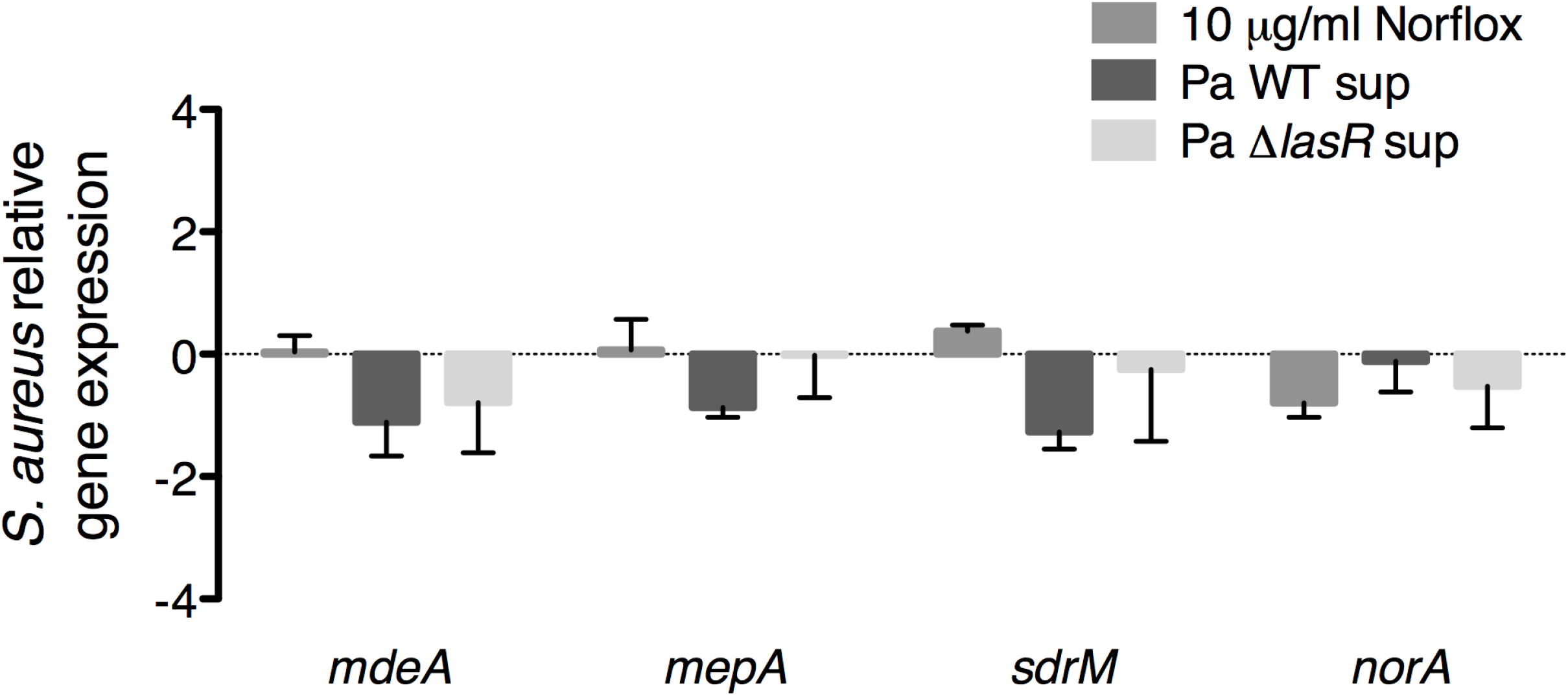
Exposure to *P. aeruginosa* exoproducts does not alter expression of a subset of *S. aureus* antibiotic transporters. Quantitative reverse transcription PCR (qRT-PCR) assays were performed to measure the expression of four *S. aureus* antibiotic transporters (*mdeA*, *mepA*, *sdrM*, *norA*) in *S. aureus* Newman biofilm populations exposed to norfloxacin (Norflox) at 10 μg/ml, supernatants from wild-type *P. aeruginosa* PA14 supernatant (Pa WT sup) and the Δ*lasR* deletion mutant (Pa Δ*lasR* sup), or medium alone. Expression was normalized to *S. aureus rpoB* and is presented as relative to expression in medium alone. Each column displays the average of three biological replicates, each with three technical replicates. Error bars indicate SD. None of the above treatment conditions resulted in significant differences in expression for any of the genes tested relative to expression in medium alone by multiple Student t tests and the Holm-Šidák method for correcting multiple comparisons.

### Testing the contribution of individual antibiotic transporters to the sensitivity of *S. aureus* biofilms to norfloxacin

Multiple *S. aureus* efflux pumps are known to transport norfloxacin; however, we wanted to rule out the possibility that a single transporter was sufficient for conferring norfloxacin resistance under our experimental conditions. Thus, we tested whether *S. aureus* strains with transposon insertions in individual antibiotic transporter genes (listed in **Table 1**) were hypersensitive to the antibiotic. Treatment with norfloxacin at either concentration tested (12.5 μg/ml or 25 μg/ml) did not significantly reduce the biofilm cell viability of the parental strain (JE2 WT) or any of the transposon mutants (**Fig. 8**). Thus, it is likely that multiple antibiotic transporters contribute to the resistance of *S. aureus* biofilms to norfloxacin under our conditions.

**Figure 8.**
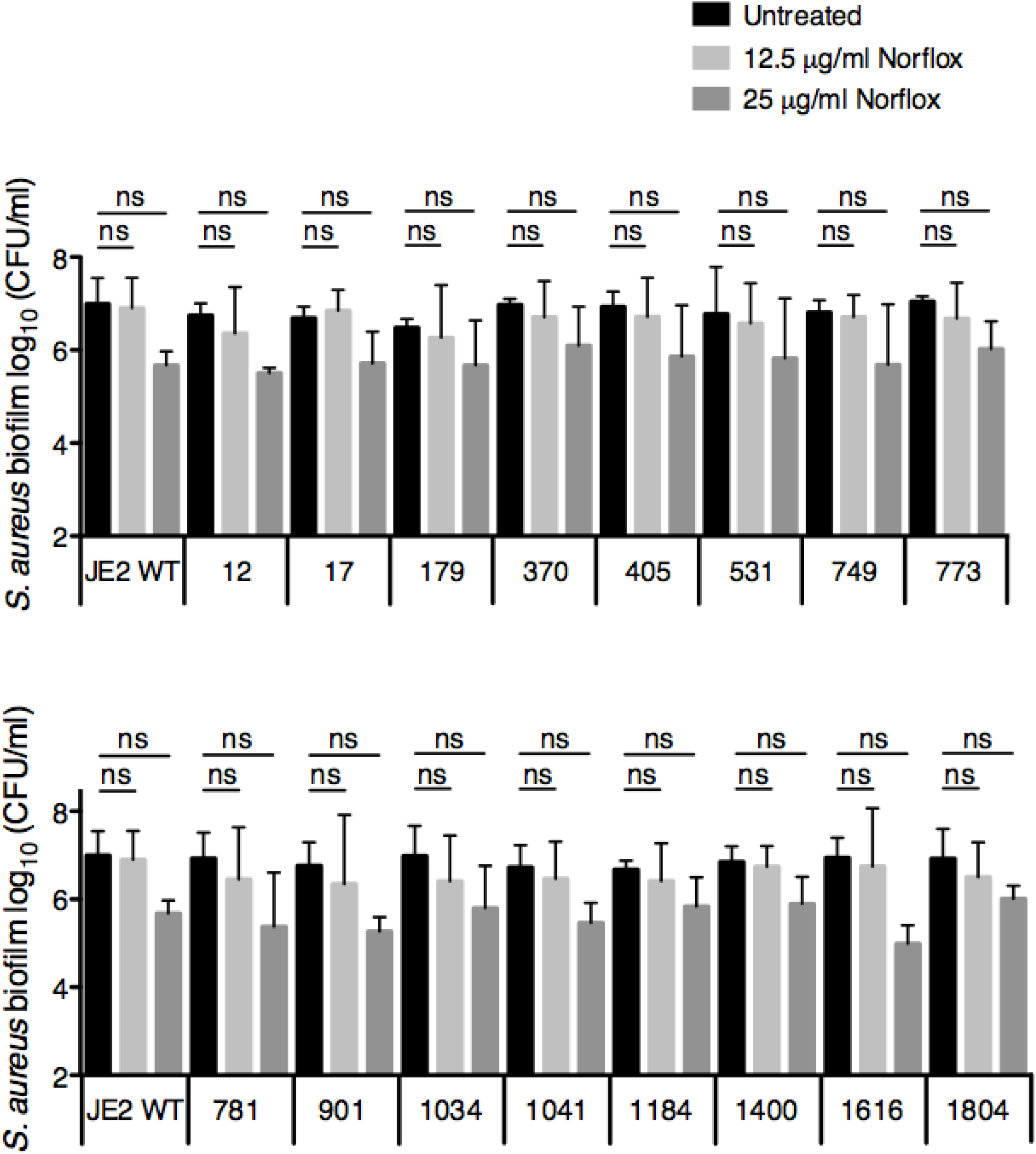
Testing the susceptibility of biofilm-grown *S. aureus* antibiotic transporter mutants to norfloxacin. **(A and B)** Biofilm disruption assays on plastic were performed with *S. aureus* JE2 parental strain or the specified transposon mutant (identified by the Nebraska transposon mutant library number, and described in **Table 1**), and norfloxacin (Norflox) at 12.5 μg/ml or 25 μg/ml. Biofilms were grown for 6 hours, exposed to the above treatments for 18 hours, and *S. aureus* biofilm CFU were determined. Each column displays the average of two biological replicates, each with three technical replicates. Error bars indicate SD. ns, not significant; by ordinary one-way ANOVA and Bonferroni multiple comparison post-test.

## Discussion

In this study, we discovered that *P. aeruginosa* influences the anti-staphylococcal efficacy of the fluoroquinolone norfloxacin. Alone, norfloxacin at concentrations of 12.5 or 25 μg/ml has minimal efficacy against *S. aureus* biofilms under our study conditions. Strikingly, we observed that *P. aeruginosa* cell-free culture supernatant increased the efficacy of this antibiotic against early (6 h) and more mature (24 h) *S. aureus* biofilms by 3-4 logs. We found multiple LasR-regulated exoproducts, including HQNO, siderophores, rhamnolipids, and phenazines likely contribute to the ability of *P. aeruginosa* to shift the susceptibility of *S. aureus* biofilms to norfloxacin.

A potential mechanism by which *P. aeruginosa* may be influencing the susceptibility of *S. aureus* to norfloxacin is via an increase in intracellular drug levels within *S. aureus* cells. Indeed, we observed that exposure to *P. aeruginosa*-secreted products led to an increase in *S. aureus* norfloxacin accumulation. Increased drug accumulation may be a consequence of increased drug uptake or decreased efflux, or a combination of the two – the data presented here suggests that both mechanisms might be at work.

In support of a model of increased influx of this antibiotic upon exposure of *S. aureus* to *P. aeruginosa* exoproducts are the following observations. Exposure to the known fluidizing agent benzyl alcohol led to a modest increase in drug efficacy, while the addition of exogenous unsaturated fatty acids (which enhance membrane fluidity) (39, 40) increased norfloxacin’s ability to kill *S. aureus* biofilms by multiple logs. Furthermore, *P. aeruginosa*-secreted factors HQNO and rhamnolipids have both been reported to alter multiple properties of the *S. aureus* cell membrane, including membrane potential, fluidity and permeability (28, 29). Thus, it is possible that these exoproducts facilitate increased diffusion of norfloxacin across the *S. aureus* cell membrane.

Additionally, several of our findings support a model of decreased efflux. In particular, we previously observed that *P. aeruginosa*-secreted products can dissipate *S. aureus* membrane potential (28). Here, we observed that treatment with the PMF inhibitor CCCP, which is well known to inhibit the activity of *S. aureus* efflux pumps (45), increased *S. aureus* intracellular norfloxacin levels and enhanced the anti-staphylococcal efficacy of norfloxacin under our study conditions, recapitulating the effect of *P. aeruginosa* supernatant on these phenotypes. Together, these data suggest the possibility that *P. aeruginosa* exoproducts interfere with the ability of *S. aureus* to effectively efflux the antibiotic. Future work is needed to elucidate the precise mechanisms and players involved in this interspecies interaction.

Overall, our results underscore the impact of polymicrobial interactions on antimicrobial efficacy and highlight the wide-ranging effects of *P. aeruginosa* exoproducts on *S. aureus* physiology. As we search for new strategies to combat recalcitrant infections caused by biofilm-based, polymicrobial infections we must consider the impact of this particular environment on the antibiotic discovery pipeline. That is, most antibiotics are developed and tested in the context of single microbes, but efficacy in such systems does not always translate in polymicrobial settings. For the example presented here, norfloxacin, but not the other quinolone antibiotics tested, showed robust activity versus *S. aureus* biofilms in the presence of *P. aeruginosa* exoproducts; this is not an observation one would predict or expect based on the activity of these antibiotics versus pure cultures. Thus, in addition to using polymicrobial models to potentially identify new antimicrobial strategies, such communities could serve as “testing platforms” to identify efficacious antibiotics *in vitro* and thus guide therapeutic choices *in vivo.*

## Materials and Methods

### Bacterial strains and culture conditions

The *S. aureus* and *P. aeruginosa* strains used in this study are listed in **Table 2**. *S. aureus* was grown in tryptic soy broth (TSB) and *P. aeruginosa* was grown in lysogeny broth (LB). Overnight cultures were grown shaking at 37°C for 12-14 h.

**Table 2.**
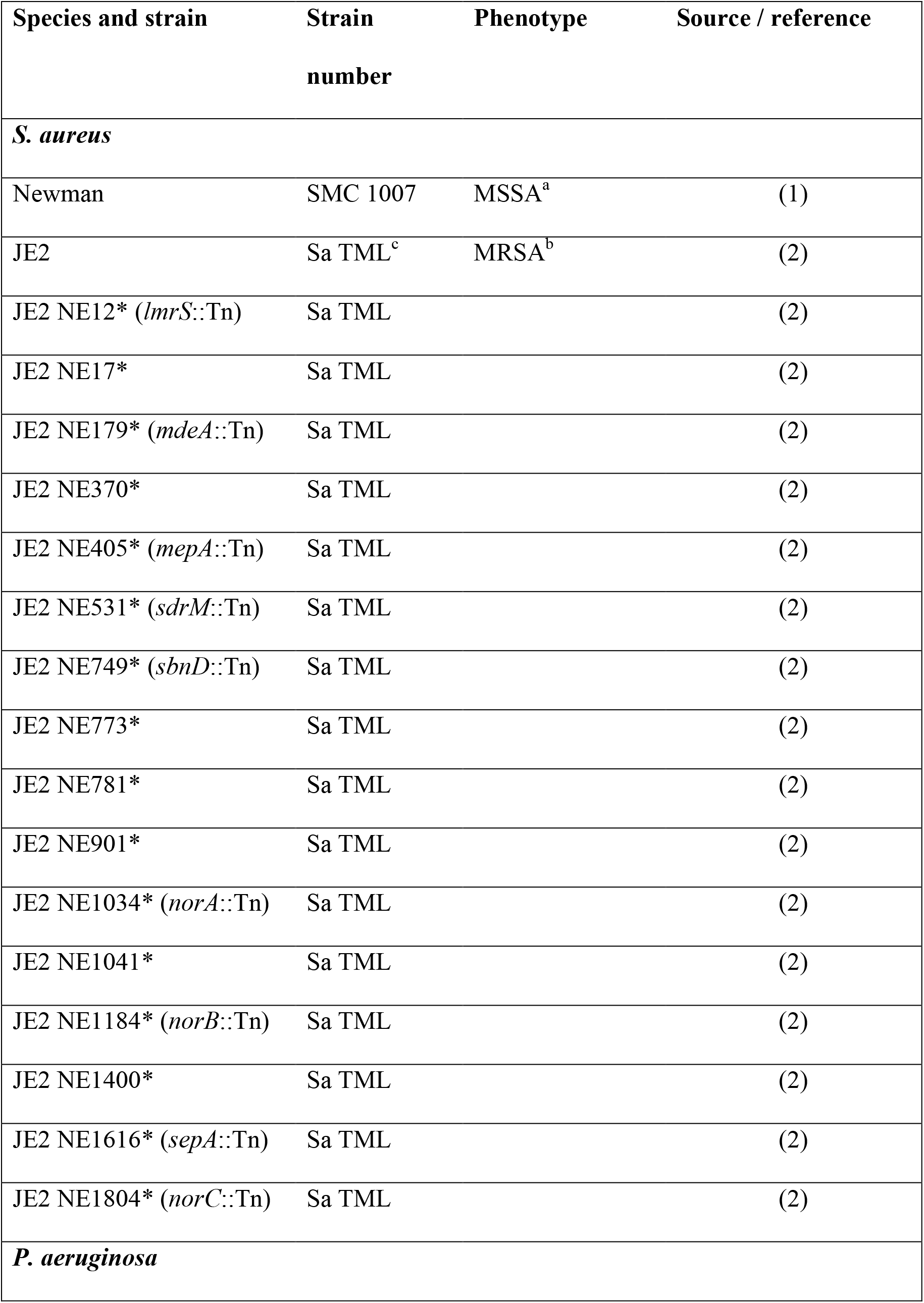
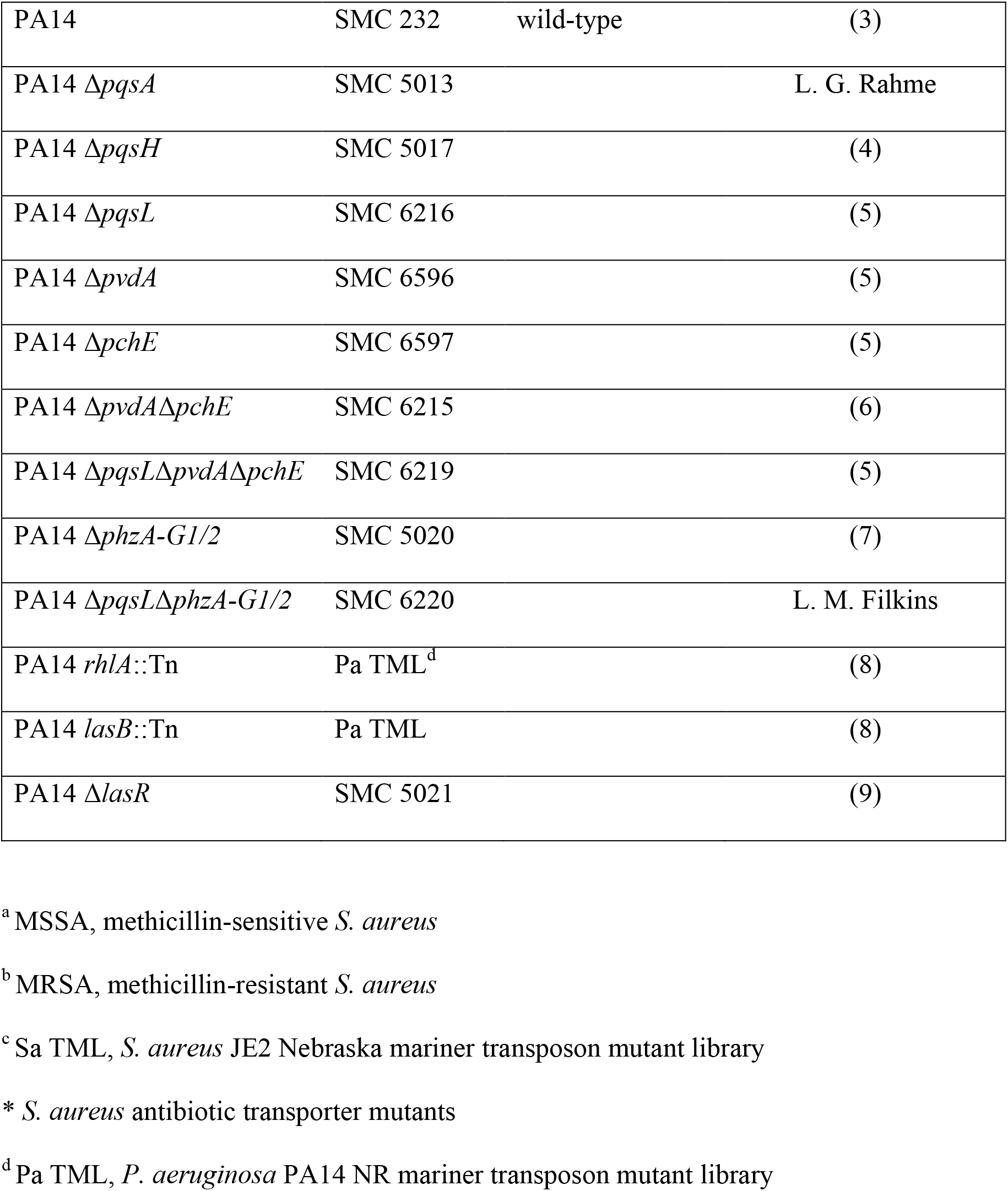
Strains used in this study.

### Preparation of *P. aeruginosa* supernatants

Overnight liquid cultures of *P. aeruginosa* were diluted to an OD_600_ of 0.05, washed in phosphate-buffered saline (PBS), and resuspended in minimum essential medium (MEM, Thermo Fisher Scientific) supplemented with 2 mM L-glutamine (MEM + L-Gln). Next, each well of a plastic 6-well plate was inoculated with 2 ml of the *P. aeruginosa* suspensions and incubated at 37°C, 5% CO_2_ for 1 h. Afterwards, unattached cells were removed and 2 ml of MEM + L-Gln was added to each well. Following incubation for an additional 22-24 h at 37°C, 5% CO_2_, the culture supernatant were collected, centrifuged at 5,000 x *g* for 5 min, and passed through a 0.22-μm filter. In subsequent experiments, wells containing *P. aeruginosa* supernatant received half the well volume of supernatant and half the well volume of either MEM + L-Gln or antibiotic solutions in MEM + L-Gln.

### Biofilm disruption assay on plastic

Overnight liquid cultures of *S. aureus* were diluted to an OD_600_ of 0.05, washed in PBS, and resuspended in minimum essential medium (MEM, Thermo Fisher Scientific) supplemented with 2 mM L-glutamine (MEM + L-Gln). Triplicate wells of a plastic 96-well plate were inoculated with 100 μl of the *S. aureus* suspensions and incubated at 37°C, 5% CO_2_. Unattached cells were removed 6 h p.i. or 24 h p.i. (as indicated), at which point antibiotic dilutions in MEM + L-Gln, *P. aeruginosa* supernatant, and MEM + L-Gln were added (total well volume of 90 μl). The plate was incubated at 37°C, 5% CO_2_ for 18 additional h. Planktonic cells were removed, and then 50 μl of PBS was added to each well of the plate. Next, the wells were scraped using a solid multi-pin replicator to disrupt the biofilms. Biofilm cells were serially diluted 10-fold in PBS, and plated on mannitol salt agar. Viable cell counts for all relevant experiments are reported as log_10_ transformed CFU/ml.

### Norfloxacin accumulation assay

Antibiotic internalization assays were performed as previously described (49–52) with modifications. Overnight liquid cultures of *S. aureus* were diluted to an OD_600_ of 2.0, washed in PBS, and resuspended in minimum essential medium (MEM, Thermo Fisher Scientific) supplemented with 2 mM L-glutamine (MEM + L-Gln). Next, 1 ml of the *S. aureus* suspension was mixed with 1 ml of either MEM + L-Gln, *P. aeruginosa* supernatant, or CCCP (for a final *S. aureus* OD_600_ of 1.0), then added to a 6-well plate (total well volume of 2 ml). The plate was incubated at 37°C, 5% CO_2_ statically for 3 h. Cells were collected, washed once with 1 ml of PBS, and adjusted to an OD_600_ of 1.0. Then, cells were resuspended in 1 ml of either MEM + L-Gln, *P. aeruginosa* supernatant, or CCCP, and added to 1 ml of PBS containing a final concentration of norfloxacin of 10 μg/ml. Afterwards, cells were incubated at 37°C under agitation for 30 min. Cells were collected and washed once with 1 ml of PBS. Next, cells were resuspended in 2 ml of 0.1 M Glycine-HCl buffer (pH of 3), followed by measurement of OD_600_ and incubation at room temperature under agitation overnight (~15 h). After centrifugation at 10,000 x g for 5 min, supernatants were transferred to a new tube. For each sample, 150 μl of supernatant was transferred to each well of clear flat-bottomed black 96-well plates. The intrinsic fluorescence of norfloxacin was measured at 440 nm following excitation at 281 nm. A standard curve was used to relate fluorescence values to the amount of intracellular norfloxacin (ng) in each sample. Values are reported as the amount of intracellular norfloxacin (ng) / OD_600_.

### RNA extraction

Overnight liquid cultures of *S. aureus* were diluted to an OD_600_ of 0.05, washed in PBS, and resuspended in minimum essential medium (MEM, Thermo Fisher Scientific) supplemented with 2 mM L-glutamine (MEM + L-Gln). Duplicate 6-well plates per condition were inoculated with the *S. aureus* suspensions (total well volume of 2 ml) and incubated at 37°C, 5% CO_2_. Unattached cells were removed at 24 h p.i., at which point antibiotic dilutions in MEM + L-Gln, *P. aeruginosa* supernatant, and MEM + L-Gln were added (total well volume of 2 ml). The plate was incubated at 37°C, 5% CO_2_ for 3 additional h. Supernatants were discarded, and then 300 μl of RNA *later* (Qiagen) was added to each well of the plate. Biofilm cells were removed from the bottom of each well using plastic cell scrapers and centrifuged at 6000 x *g* for 5 min at 4°C. Following removal of supernatants, total RNA was isolated using PureZOL RNA Isolation Reagent (BioRad) according to the manufacturer’s instructions, including homogenization by bead beating (5 cycles of 30 sec). Afterwards, Turbo DNA-*free* DNase treatment (Thermo Fisher Scientific) was performed according to the manufacturer’s instructions followed by PCR to test for DNA contamination.

### Quantitative reverse transcription PCR

cDNA synthesis and quantitative reverse transcription PCR (qRT-PCR) were performed using the iTaq Universal SYBR Green One-Step Kit (BioRad) according to the manufacturer’s instructions. The qRT-PCR primers used were designed for each *S. aureus* gene of interest and are listed in **Table 3**. *S. aureus rpoB* was selected to normalize gene expression as determined previously (30). For each treatment condition, cycle threshold (C_T_) values for each gene of interest were normalized to *rpoB* C_T_ values. Changes in gene expression were assessed by the delta-delta C_T_ method (53). Expression data are presented as *rpoB*-normalized gene expression for each treatment condition relative to *S. aureus* exposed to medium alone.

**Table 3.**
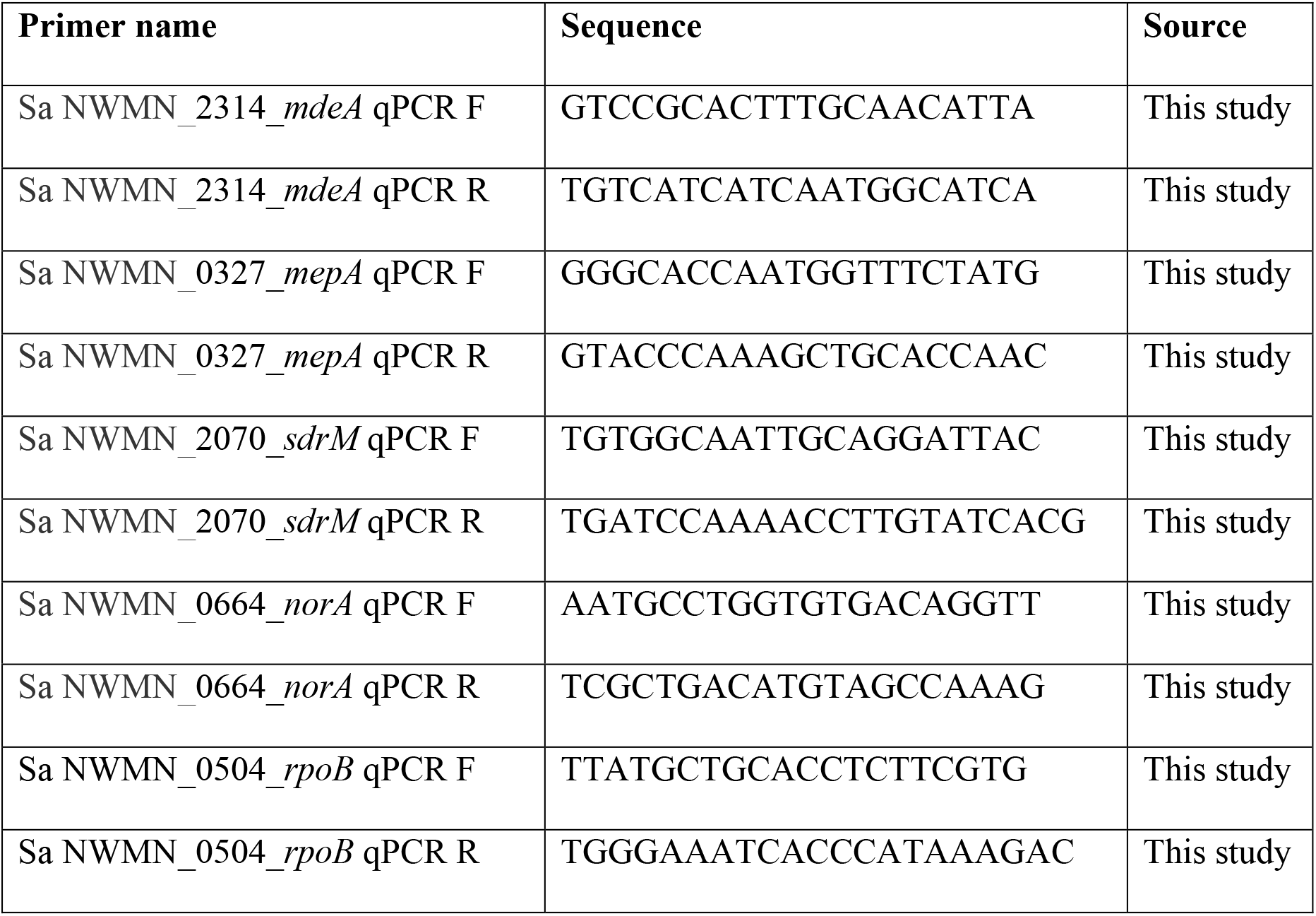
Primers used in this study.

## Acknowledgements

We thank Ambrose Cheung, Laura Filkins, and Deborah Hogan for providing bacterial strains.

